# Comparing cancer cell lines and tumor samples by genomic profiles

**DOI:** 10.1101/028159

**Authors:** Rileen Sinha, Nikolaus Schultz, Chris Sander

## Abstract

Cancer cell lines are often used in laboratory experiments as models of tumors, although they can have substantially different genetic and epigenetic profiles compared to tumors. We have developed a general computational method – TumorComparer - to systematically quantify similarities and differences between tumor material when detailed genetic and molecular profiles are available. The comparisons can be flexibly tailored to a particular biological question by placing a higher weight on functional alterations of interest (‘weighted similarity’). In a first pan-cancer application, we have compared 260 cell lines from the Cancer Cell Line Encyclopaedia (CCLE) and 1914 tumors of six different cancer types from The Cancer Genome Atlas (TCGA), using weights to emphasize genomic alterations that frequently recur in tumors. We report the potential suitability of particular cell lines as tumor models and identify apparently unsuitable outlier cell lines, some of which are in wide use, for each of the six cancer types. In future, this weighted similarity method may be generalized for use in a clinical setting to compare patient profiles consisting of genomic patterns combined with clinical attributes, such as diagnosis, treatment and response to therapy.

## Introduction

Immortalized cancer cell lines, derived from tumors and grown and maintained *in vitro,* are the most commonly used experimental models in cancer research. Cell lines preserve many properties of tumors, and have been of immense value in advancing the understanding of cancer biology and developing novel therapies over the past decades ^1-3^. However, there are important differences, both in general and in particular tumor types, between the genetic alteration profiles of cell lines and tumors, which are the subject of this study.

While cell lines retain many features of tumors, they also acquire additional changes in the process of immortalization, and during growth and maintenance in culture. Several studies have reported differences between cell lines and tumors with respect to gene expression ^4,5^, methylation ^6^–^8^, and copy number alterations ^9,10^. In general, cell lines tend to have more genomic alterations than primary tumors, which can be explained by a bias towards using cell lines derived from metastatic tumors ^1^, and *in-vitro* selection of subpopulations of cell lines during long periods of growth and maintenance in the laboratory ^1^. Furthermore, the apparent overall difference in mutation burden between cell lines and tumors may be affected by the presence of germline mutations in cell lines, which are explicitly removed from tumor data as matched normal samples are usually available for tumors, but which are incompletely removed from cell line data even with the customary filtering of known common germline variants. In addition, as there is a systematic bias in the source of most immortalized cell lines, cell lines typically do not represent all subtypes of cancers in a particular tissue of origin. In particular, tumor subtypes with the least amount of genetic alterations tend to be under-represented ^11-13^. Given these differences, selecting the most suitable cell line(s) for a specific laboratory investigation becomes a technical challenge of practical interest. In general, cell lines with profiles similar to tumor samples are more suitable than outliers. However, when a set of particular features, such as mutations in particular oncogenes, are required for cell lines to “phenocopy” aspects of tumors, focus on these features would provide more useful assessment of similarity^14^.

Thus, beyond overall genetic similarity, the choice of an appropriate cell line for a specific scientific project crucially depends on the goal and context of the study and comparison algorithms should take the investigator’s interest into account. For example, one may want to choose a cell line that is most similar to a set of tumors in terms of alterations in signalling pathways, such as protein phosphorylation cascades; or, in terms of mutations in particular pathways; or, in terms of the overall level of alterations in known oncogenic pathways.

We therefore aimed to develop a general method for adapting the criteria for the choice of cell line most similar to a particular tumor type to the biological question at hand. A simple yet powerful approach is to incorporate feature weights into the measure of similarity of molecular profiles. For example, alterations in genes involved in a certain type of signalling may get a higher weight, others a lower weight. A very simple choice of weights is 1.0 (chosen) and 0.0 (ignored), but in general weights are real numbers 0.0≤w≤1.0. Here, we aimed to derive weights that emphasize potentially oncogenic genomic alterations, while de-emphasizing alterations that are likely to be “passengers” in tumors. We derived such weights (called RA1, for TumorComparer weights based on Recurrent Alterations 1) from TCGA tumor profiles, and then computed the weighted similarity between tumors and cell lines using these weights. We applied the method with RA1 weights to compare tumors of 6 different cancer types from The Cancer Genome Atlas (TCGA) ^15^ to cell lines from the Cancer Cell Line Encyclopedia (CCLE) ^16^, and identified good, moderate and poor matches as well as outlier cell lines to guide cell line selection for laboratory experiments focussed on oncogenic processes.

## Results

We developed an approach to comparing tumors and cell lines using multiple data types, using weights to emphasize events relevant to the biological question at hand, e.g. the more frequent and/or known oncogenic events (Figure 1). We then applied our method to compare cell lines and tumors for six different cancer types, using genomic data from 260 CCLE cell lines and 1914 TCGA tumor samples (Table 1) and weights emphasizing recurrent alterations in tumors. By investigating the nearest neighbours of the cell lines and tumors, we identified the best matching cell lines for the tumors of various types, as well as poor matches and outlier cell lines.

**Figure 1.**
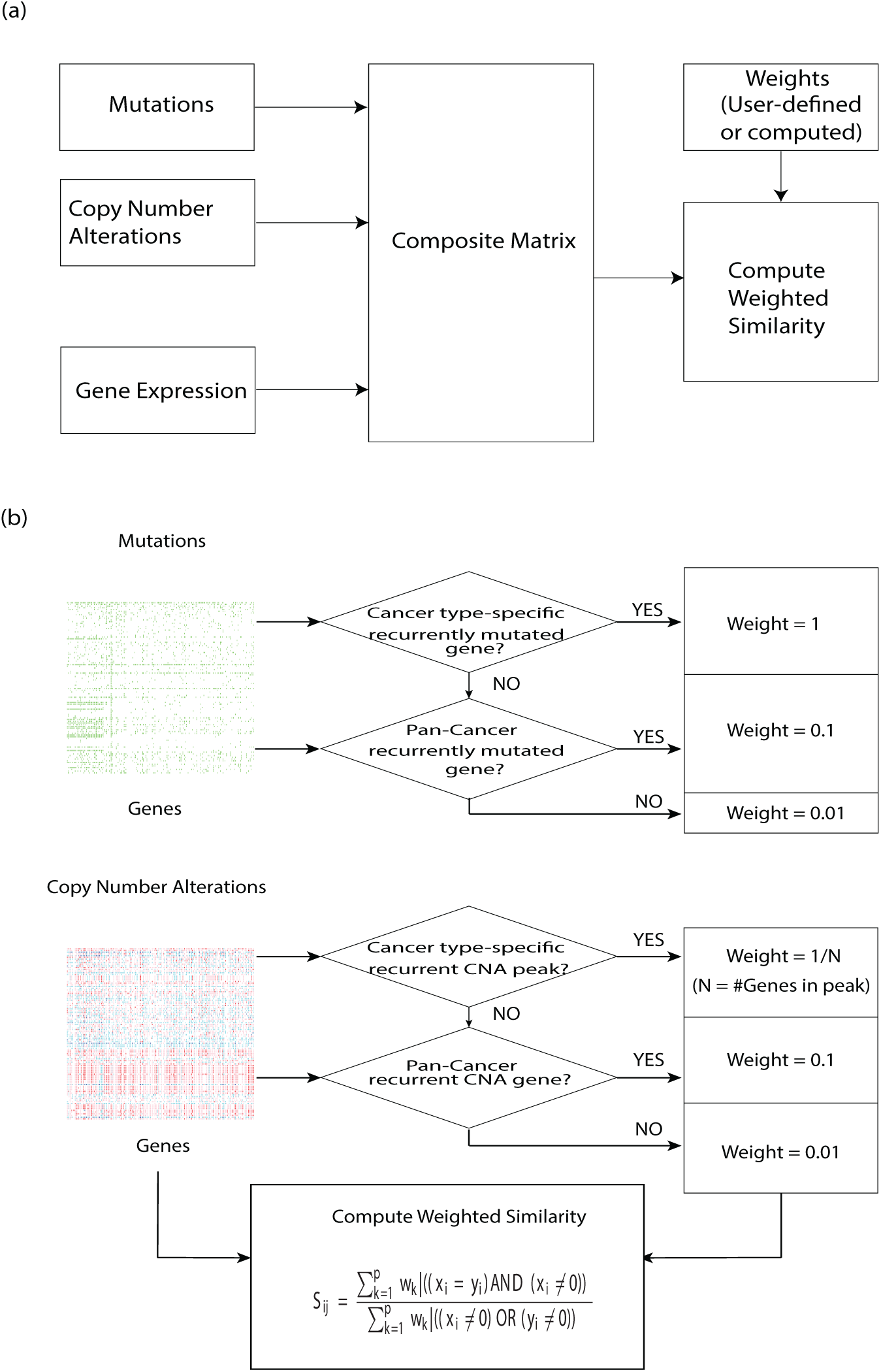
Workflow of the TumorComparer. (a) Weighted similarity is computed using weights that are either derived from data or provided by the user, and reflect the emphasis placed on particular genetic properties of tumors. Multiple data types, such as mutations and DNA copy number alterations, are combined into one composite matrix containing all features to be compared. (b) To compare cell lines and tumors, we used mutations (mutated - green, wild type - white) and copy number alterations (gains – light and dark red, losses – light and dark blue, diploid - white), and chose weights based on recurrence of cancer-type specific and/or pan-cancer events, as a proxy for likely functional events.

**Table 1.** Tumor and cell line datasets. The 1914 tumors from The Cancer Genome Atlas (TCGA) and 260 cell lines from the Cell Line Encyclopedia (CCLE) used here represent 6 cancer types/subtypes.

| Cancer Type | # of Tumors | # of Cell Lines |
| --- | --- | --- |
| Ovarian Cancer (OV) | 311 | 47 |
| Lung Squamous Cell Carcinoma (LUSC) | 178 | 25 |
| Lung Adenocarcinoma (LUAD) | 172 | 42 |
| Breast Cancer (BRCA) | 760 | 51 |
| Colorectal Adenocarcinoma (COADREAD) | 220 | 54 |
| Glioblastoma (GBM) | 273 | 41 |

### Tumors vary more in the extent of alterations by cancer type than do cell lines

We compared mutations and copy number alterations (CNAs) in cell lines from the Cancer Cell Line Encyclopedia (CCLE) and tumors from The Cancer Genome Atlas (TCGA) for six different cancer types (lung adenocarcinoma - LUAD, lung squamous cell carcinoma - LUSC, high grade serous ovarian carcinoma - OV, breast carcinoma - BRCA, colorectal adenocarcinoma - COADREAD, and glioblastoma - GBM). We used cell lines of the corresponding types/subtypes, with two exceptions – all ovarian cell lines were used (since the type annotation is not always unambiguous, and is debatable in some cases, as we recently reported ^17^), and all CCLE cell lines annotated as “large intestine” were used as colorectal cell lines.

CCLE provides mutation data for 1651 genes, and we restricted the current analysis to using CNA data for the same subset of genes. CNA data was available for 1529 of these 1651 genes, giving us 3180 alterations or features altogether. Mutations were represented in binary form (1 = ‘gene is mutated’, 0-‘gene is not mutated’), irrespective of location and number of mutations. For copy number changes we used a 5-valued (-2,-1,0,1,2) GISTIC ^18^ representation.

While our main focus is on detailed comparison of individual tumor profiles with those of potentially useful cell line models, some general trends emerge from the systematic comparison across many cell lines and tumors in a number of different cancer types. First, we confirm the previous observation ^6,9,13,17^ that cell lines tend to have a larger number of genes affected by somatic mutations or copy number alterations than do tumors (Figure 2). While the lack of matched germline samples for cell lines may confound this conclusion, it plausibly remains valid because of the systematic removal of common variants from the set of non-synonymous mutations in the CCLE cell line dataset. The higher level of genetic alterations in cell lines may simply be the consequence of their origin in metastatic cancers or of their adaptation to laboratory conditions, as well as their higher purity when compared to tumor samples. Whether or not cell lines are nonetheless good models for certain tumor types depends on the extent to which they retain alterations characteristic of human tumor tissue, which is analysed in detail below.

The second general observation relates to the extent of genetic variation in tumors and cell lines derived from different tissues. We confirm the observation that mutation counts, as well as copy number alterations, are substantially higher for some tumor types than others (Figure 2) ^19,20^. In contrast, cell lines vary less between different tumor types of origin (Figure 2). This observation may be related to mechanisms of immortalization, in-vitro growth or adaptation via passaging.

**Figure 2.**
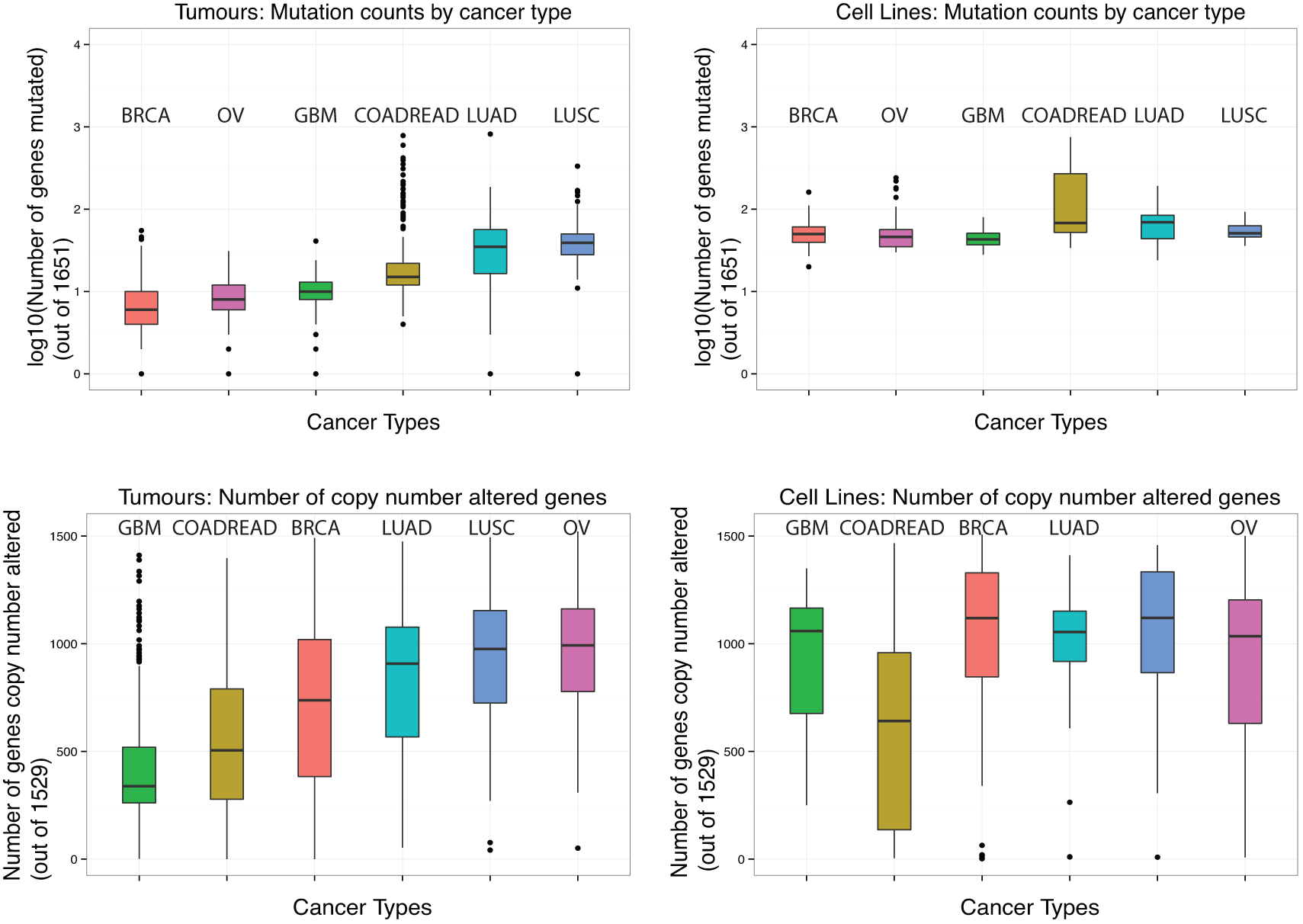
Tumors vary more in the extent of mutations and copy number alterations by cancer type than do cell lines. Comparison of the extent of mutation (top) and copy number alteration (CNA, bottom) across six cancer types for cell lines and tumors. While tumors vary remarkably in mutation counts (top left; e.g., the median number of mutations in lung squamous cell carcinoma is nearly six times greater than in breast cancer and five times greater than in high grade serous ovarian carcinoma), the corresponding variation is less than two-fold in cell lines (top right). The trend for copy number alterations (bottom) is similar, albeit less pronounced. This difference between tumors and cell lines may be due to alterations acquired by cell lines during immortalization, in-vitro growth and/or adaptation via passaging.

The trend for copy number alterations (bottom) is similar, albeit less pronounced. This difference between tumors and cell lines may be due to alterations acquired by cell lines during immortalization, in-vitro growth and adaptation via passaging.

### All cancer types have a few good tumor matches, and most have some outlier cell lines

To ensure that known and likely oncogenic genomic alterations are emphasized in the comparisons, while differences in likely insignificant or “passenger” alterations are de-emphasized, we aimed to select weights that reflect oncogenic events in tumors. To this end, we utilized the results of the MUTSIG ^19,21^ and GISTIC ^18^ programs for TCGA mutation and CNA data, respectively, and gave higher weights to genes that were identified by MUTSIG as being significantly recurrently mutated, reported by GISTIC as being in a significant CNA peak, and intermediate weights to other genes known to be important in cancer (based on the TCGA pan-cancer analyses of mutation ^22^ and CNA data ^20^). All other genes had a lower, default weight (see Methods for details).

We assessed the suitability of each cell line as a tumor model using its weighted similarity to the tumors (weight set RA1), calculated as the weighted asymmetric matching score, ignoring zero-zero matches (see Methods for details). Since tumors are themselves a heterogeneous group, often consisting of multiple subtypes, we looked at the mean similarity to *k* nearest tumors (MSK), instead of mean similarity to the entire tumor cohort. Results are shown for k=10% of tumors in the respective dataset; results were not significantly different for other values of k, e.g. 20%, 30% etc.). We also compared the MSK values of tumors, which estimate tumor-tumor similarity, to the MSK value of each cell line to ascertain whether a given cell line is a good genomic match, a moderately good match, a poor match or an outlier (Table 2).

For approximate visual assessment of the proximity of tumors and cell lines, we project them into two dimensions (Figure 3) using multidimensional scaling (MDS) with all-against-all distances between samples as input. Intuitively, cell lines that are close to several tumors on the MDS plots are good genomic matches, those which are far from most tumors are poor matches and outliers, while the remaining cell lines are intermediate matches. Figure 4 shows the MSK scores for all 260 cell lines across the 6 tumor types – the number of cell lines which are good matches (near or exceeding the mean tumor-tumor similarity, near or above the green line) or particularly poor matches (more than two standard deviations away from the mean tumor-tumor similarity levels; below the red line).

**Figure 3.**
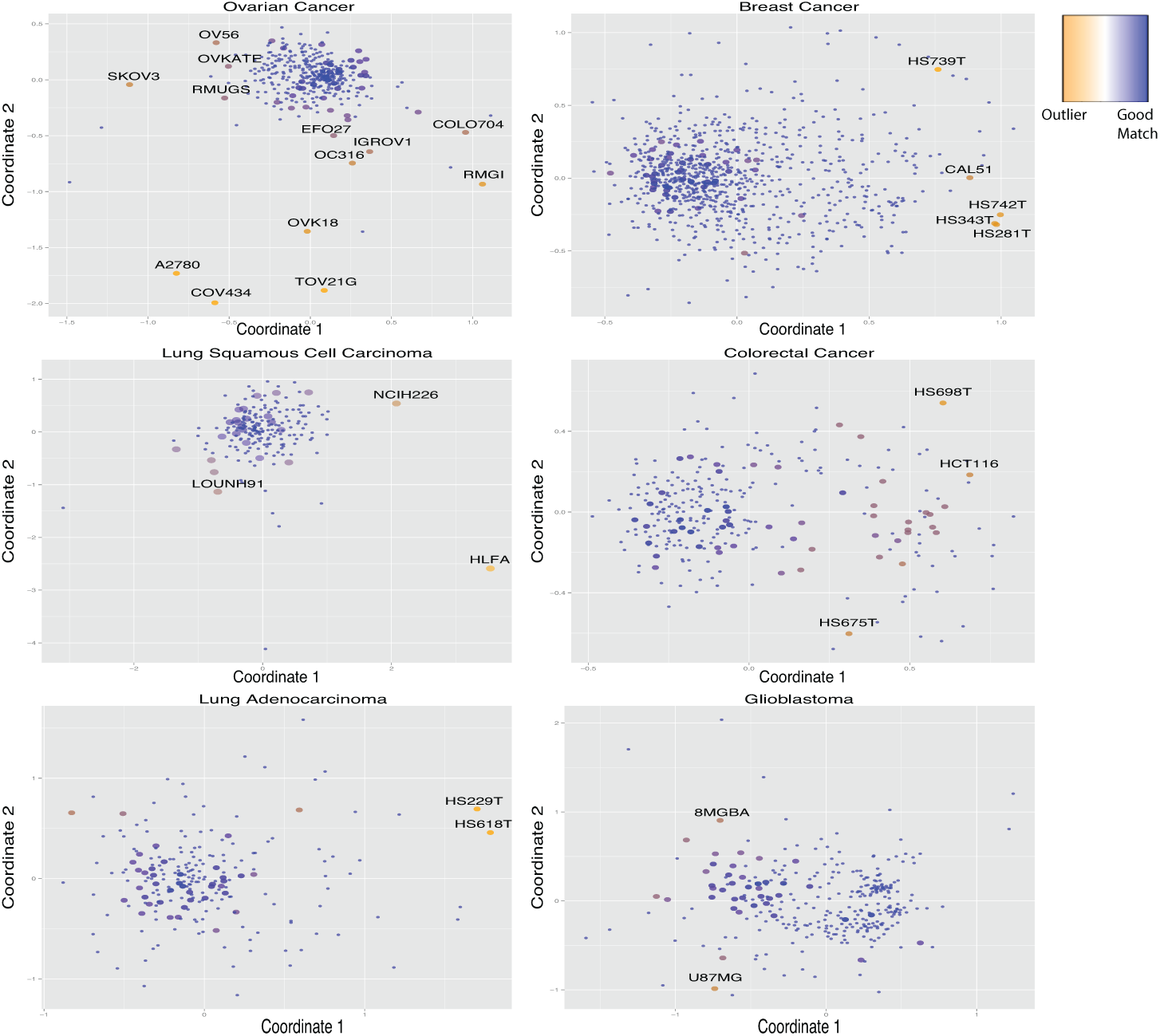
All six cancer types have cell lines that are good tumor matches and some outlier cell lines. The spread of the tumors shows that OV and LUSC are more homogenous than BRCA and LUAD. The position of cell lines in the two-dimensional representation (multi-dimensional scaling) relative to the tumors indicates which cell lines are good matches (near some tumors) and which are poor matches or outliers (far from most/all tumors). Similarities between tumors (small blue dots) from TCGA tumor (sub)types and corresponding cell lines (larger dots, orange to blue, from outliers to good matches) from CCLE were computed using mutation and copy number alteration data with weights reflecting cancer-specific significant genomic alterations (RA1). In the interest of clarity, only the outlier cell lines are labelled (further details in Methods and Supplement Table 1).

**Figure 4.**
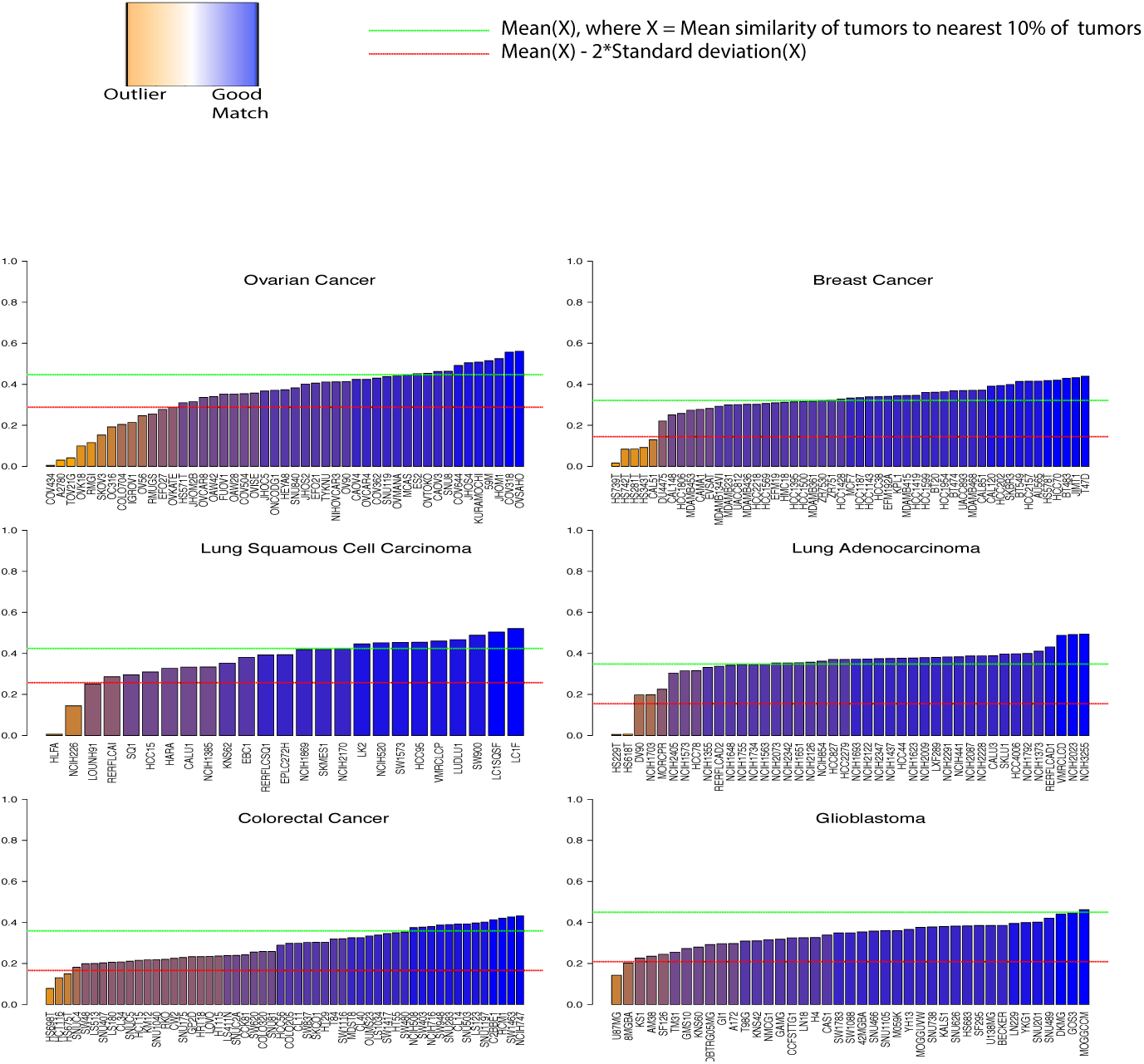
Most cell lines are moderately good matches to tumors, but some cell lines are clear outliers. Cell line MSK scores (mean similarity to k nearest tumors), with tumor MSK scores in the foreground for six cancer types (k = 10% of tumor dataset, green line: mean tumor MSK score, red line: mean (tumor MSK scores) – 2*sd(tumor MSK scores), the threshold for outliers). The MSK scores show that there is a spectrum of good and moderately good to poor tumor matches and outlier cell lines.

### Several ovarian cell lines are poor matches to TCGA high-grade serous ovarian tumors; A2780, SKOV3 and IGROV1 are among the highly cited outliers

High grade serous ovarian carcinoma (HGSOC) is characterized by a relatively low number of mutations (but near-ubiquitous TP53 mutations), and a medium to high extent of copy number alterations ^23^. We recently evaluated 47 ovarian cell lines from the CCLE as models of HGSOC using the genomic profiles of TCGA tumors, and found that several highly cited cell lines are poor genomic matches to the tumors, while some less popular cell lines make for better matches ^17^. Here, we revisited the comparison of ovarian cell lines and tumors using the weighted similarity approach.

On average the MSK for HGSOC tumors was 0.45. Eleven cell lines (ES2, OVTOKO, CAOV3, SNU8, COV644, JHOS4, KURAMOCHI, 59M, JHOM1, COV318 and OVSAHO) scored higher than this (Figure 4), indicating that they might be particularly good matches with respect to the genomic alterations we chose to emphasize. On the other hand, COV434, A2780, TOV21G, OVK18, RMGI, SKOV3, OC316, IGROV1, COLO704, OV56, RMUGS, EFO27 and OVKATE were clear outliers, with an MSK ≤ 0.29. As noted earlier ^17^, IGROV1, OC316, EFO27, OVK18 and TOV21G have few CNAs and an exceptionally large number of mutations - the exact opposite of the majority of TGCA HGSOC tumors, which have few mutations and a medium to high level of copy number aberrations, relative to other cancer types.

The combined CCLE and TCGA dataset contained 3 mutations and 28 genes in focal CNA peaks, including 13 in singleton peaks (i.e., only one nominated target gene in the focal peak) declared significant by MUTSIG and GISTIC, respectively. Most cell lines contained some of these 16 alterations, as well as alterations in other cancer genes found in the TCGA pan-cancer studies ^20,22^. However, several of the outlier cell lines lacked the HGSOC-specific alterations, but had alterations in other cancer genes. More specifically, A2780, COV644, MCAS, OC316, OVTOKO and SNU840 lack any of the HGSOC-specific important alterations. Interestingly, the cell line JHOM2B, which has mutations in TP53 and RB1 but lacks any high level amplifications or homozygous deletions characteristic of HGSOC, also carries a BRAFV600E mutation. When all CCLE cell lines were clustered using mRNA expression (not shown), JHOM2B clustered with colorectal cell lines, raising the possibility that it might be a colorectal cell line, though further study is needed to confirm or rule that out.

### Most breast cancer cell lines, including all highly cited cell lines, are at least moderately good matches to a subset of tumors

The citation landscape of BRCA cell lines is dominated by a handful of cell lines, including MCF7, MDAMB231, T47D, MDAMB435 and MDAMB468 ^24^. All of these cell lines were found to be good genomic matches for at least 76 tumors (10% of all BRCA tumors analysed, Figure 4). The outliers were HS739T, CAL51, HS281, HS343T, and HS742T, none of which are highly cited (in fact, CAL51, with 38 citations appears to be the only one used commonly – HS742T has only one citation while HS739T, HS281T and HS343T had no PubMed hits). All five outlier cell lines have very flat CNA profiles, with almost no copy number alterations. HS281T and HS731T have very few BRCA-specific recurrent mutations. Although the five outlier cell lines are all triple-negative breast cancer (TNBC) cell lines^25^, that does not explain their low similarity to tumors, since several other CCLE TNBC cell lines (e.g. BT20, BT549 and MDAMB468) have much higher similarity scores. In general, BRCA cell lines resemble BRCA tumors much more than ovarian cell lines resemble HGSOC tumors - however, this is at least partly due to a greater number of BRCA-specific alterations (29 MUTSIG genes and 33 genes in GISTIC focal peaks). Moreover, the median tumor-tumor MSK was only 0.33 for TCGA breast tumors, indicating greater heterogeneity within the cohort, and “lowering the bar” for a cell line to be considered a good match to tumor genome profiles.

### Most lung adenocarcinoma cell lines are at least moderately good matches to tumor genomic profiles; HS229T and HS618T are outliers while poor matches include DV90, NCI-H1703 and MORCPR

Similar to BRCA cell lines, most LUAD cell lines are good matches to at least 10% of the tumors (17), except HS229T, HS618T, DV90, NCIH1703 and MORCPR, none of which are highly cited (Figure 4). Also similar to TCGA BRCA are the relatively high number of significant alterations (21 MUTSIG genes and 39 genes in GISTIC focal peaks) and relatively low median tumor-tumor MSK of 0.36. DV90, HS229T and HS618T are characterized by unusually flat copy number profiles. DV90 also has an unusually high number of mutations, along with NCI-H1573 and NCI-H2342 (which also has a high number of high-level CNAs). On the other hand, NCI-H2023, VMRCLCD and NCI-H3255 were the top-scoring LUAD cell lines with an MSK of 0.49, close to the highest tumor-tumor MSK of 0.51. RERFLCAD1 was also above the third quartile (0.42) of LUAD tumor-tumor MSK.

### NCI-H226, LOUNH91 and HLFA are outliers among lung squamous cell carcinoma cell lines, while SQ1 and RERFLCAI are also relatively poor matches

LUSC is characterized by a high extent of mutations and copy number alterations. LUSC tumors typically have a higher extent of CNAs than LUAD tumors. These characteristics are also found in the majority of LUSC cell lines, with the exception of HLFA, which has a remarkably flat CNA profile (fraction genes altered, FGA = 0.6%). LOUNH91 (20% FGA) and NCI-H226 (33% FGA) have relatively quiet CNA profiles, and also lack LUSC-specific recurrent alterations. However, while NCI-H226 is a relatively well-established cell line with 57 citations, LOUNH91 and HFLA appear to be used much less often (1 and 1 citations, respectively). NCI-H1385 stands out by virtue of having the highest number of high-level CNAs (21% genes, as opposed to a median of 3.6% for LUSC cell lines and 2.9% for LUSC tumors). HCC15 has an exceptionally high number of mutations (93 genes (5.6%), as opposed to a median of 51 genes (3.1%) for LUSC cell lines and 39 genes (2.3%) for LUSC tumors). Notably, as many as 40% (10/25) CCLE LUSC cell lines are reported as being TP53 wild type, in contrast to TCGA LUSC tumors in which 82% (146/178) carry TP53 mutations.

TCGA LUSC tumors have a relatively high median MSK of 0.42, and have 6 MUTSIG genes and 31 GISTIC genes in our dataset, consistent with the nature of a cancer that is driven more by CNAs than mutations ^26^. Out of the 25 CCLE LUSC cell lines we analysed, as many as 12 (SKMES1, HCC95, NCI-H1869, NCI-H2170, LK2, NCI-H520, SW1573, VMRCLCP, LUDLU1, SW900, LC1SQSF and LC1F) had an MSK ≥ 0.42, with SW900, LC1SQSF and LC1F exceeding the third quartile (0.48) of LUSC tumor-tumor MSK (Figure 4).

### U87MG and SF126 are among poor genomic matches for glioblastoma tumors; several glioma cell lines lack GBM-specific recurrent alterations

TCGA GBM tumors have 14 MUTSIG genes and 31 GISTIC genes in our dataset, and a relatively high median tumor-tumor MSK of 0.45. CCLE includes gliomas as a subset of “central nervous system” cell lines. We compared all CCLE glioma cell lines to the TCGA GBM tumors. U87MG (MSK 0.16), the most widely used GBM cell line (1578 citations), is a poor match for the genomic profile of the tumor cohort due to its atypical profile, as well as harbouring several alterations which are not recurrent in GBM but are important in other cancers (Figure 4). 8MGBA is an outlier for similar reasons, albeit one with relatively few (13) citations. Interestingly, several CCLE glioma cell lines lack GBM-specific recurrent alterations reported by TCGA ^27^. H4 has none of the GBM-specific mutations, and only has a homozygous deletion of PTEN among the GBM-specific GISTIC peaks. KNS60, MOGGCCM and MOGGUVW have EGFR and TP53, RB1 and TP53, and PTEN mutations respectively, but no recurrent CNAs. Similarly, SF295, SNU201 and YH13 lack any GBM-specific recurrent CNAs.

### Colorectal cell lines show a lot of intra-cancer type heterogeneity (as do TCGA colorectal tumors); HCT116, HS675T and HS698T are outliers among colorectal cell lines

TCGA COADREAD tumors have 19 MUTSIG genes and 18 GISTIC genes in our dataset, and have a median tumor-tumor MSK of 0.38, indicative of greater intra-cancer type heterogeneity than OV, LUSC and GBM, and in a similar range to BRCA and LUAD. This is in agreement with the TCGA colorectal study ^28^, which reported that colorectal cancers showed great variation in mutation rates, with a subset of tumors demonstrating microsatellite instability (often along with hypermethyation and MLH1 silencing) and carrying a much higher mutational burden than the majority of tumors, and another subset of hyper-mutators with yet higher number of mutations, somatic mismatch-repair and POLE mutations ^28^. Several of the CCLE colorectal cell lines (e.g. SNU1040, SNU81, CW2, HCT15, HT115, SNU175 and GP2D) have a very high mutation count (of 418-750, compared to a median of 68 for all CCLE colorectal cell lines). In the absence of information on POLE mutations, methylation profiles and microsatellite instability status, it is challenging to resolve the colorectal cell lines into hyper-mutants and others. Most colorectal cell lines have at least some recurrent COADREAD-specific alterations reported by TCGA, making them at least moderately good matches to a subset of COADREAD tumors. At first sight, HS698T (MSK 0.08), HCT116 (MSK 0.14) and HS675T (MSK 0.15) seem to be particularly poor matches for the TCGA colorectal tumors, with HS675T, CW2, LS180, GP2D, HCT15, SNUC4 and HS698T showing an unusually low fraction of copy number altered genes (5-60 genes; median for colorectal cell lines = 641.5). HCT116, which is very widely used (4613 citations) also has many alterations that are rare in colorectal cancer but recurrent in the TCGA pan-cancer dataset. HS698T and HS675T, on other hand, have no PubMed citations to date. However, several of these cell lines have recently reported as hypermutated^29^ (A total of 14 out of 54 CCLE colorectal cell lines - CCK81, GP2D, HCT116, HCT15, HT115, HT55, KM12, LOVO, LS180, LS411, RKO, SNU175, SNUC4, SW48), which would explain the high number of mutations and low extent of copy number alterations, making these cell lines representatives of the hypermutated subtype of colorectal cancer rather than outliers. Since the hypermutated subtype is characterized by an exceptionally high mutational burden rather than specific recurrent alterations, comparisons based on finding shared recurrent events (such as here with weights RA1) are likely to identify hypermutated samples as poor matches to most tumors and/or potential outliers.

**Table 2.** 28 outlier cell lines from 6 cancer types. The genomic profiles of these cell lines are badly matched to tumors from this cancer type. These cell lines are very probably not good models for tumors. The cell line HCT116 has been recently reported to be hypermutated^29^. Details of alterations for each cell line are in the supplement (Tables S1-S6).

| **Cancer Type or Subtype** | **Cell Line** | **Number of Citations** | **Main Atypical Alterations** |
| --- | --- | --- | --- |
| Breast Cancer | HS742T | 1 | Flat CNA profile |
| Breast Cancer | HS739T | 0 | Flat CNA profile, also has few BRCA-specific recurrent mutations |
| Breast Cancer | HS343T | 0 | Flat CNA profile |
| Breast Cancer | HS281T | 0 | Flat CNA profile, also has few BRCA-specific recurrent mutations |
| Breast Cancer | CAL51 | 38 | Flat CNA profile |
| Lung Adenocarcinoma | HS618T | 0 | Flat CNA profile, also lacks LUAD-specific recurrent mutations |
| Lung Adenocarcinoma | HS229T | 0 | Flat CNA profile, also lacks LUAD-specific recurrent mutations |
| Lung Squamous Cell Carcinoma | NCIH226 | 57 | Relatively quiet and atypical CNA profile; also lacks LUSC-specific recurrent mutations |
| Lung Squamous Cell Carcinoma | HLFA | 2 | Flat CNA profile, also lacks LUSC-specific recurrent mutations |
| Lung Squamous Cell Carcinoma | LOUNH91 | 1 | Relatively quiet and atypical CNA profile |
| Glioblastoma | U87MG | 1578 | Atypical CNA profile; also has several mutations and CNAs in cancer genes not typically altered in GBM |
| Glioblastoma | 8MGBA | 13 | Atypical CNA profile; also has several mutations and CNAs in cancer genes not typically altered in GBM |
| Colorectal Cancer | HS675T | 0 | Flat CNA profile; also has few COADREAD-specific recurrent mutations |
| Colorectal Cancer | HCT116 | 4613 | Flat CNA profile; also has many mutations in cancer genes not typically mutated in COADREAD. Might represent the hypermutated subtype of colorectal cancer<sup>29</sup>. |
| Colorectal Cancer | HS698T | 0 | Flat CNA profile; also has few COADREAD-specific recurrent mutations |
| Ovarian Cancer | A2780 | 1812 | Flat CNA profile |
| Ovarian Cancer | IGROV1 | 309 | Hypermutated; flat CNA profile |
| Ovarian Cancer | EFO27 | 21 | Hypermutated |
| Ovarian Cancer | COV434 | 41 | Flat CNA profile |
| Ovarian Cancer | COLO704 | 1 | Hypermutated |
| Ovarian Cancer | OVKATE | 1 |  |
| Ovarian Cancer | OVK18 | 10 | Hypermutated |
| Ovarian Cancer | OV56 | 0 | Atypical CNA profile |
| Ovarian Cancer | OC316 | 3 | Hypermutated |
| Ovarian Cancer | TOV21G | 53 | Hypermuted; flat CNA profile |
| Ovarian Cancer | SKOV3 | 2918 | Fairly quiet and atypical CNA profile |
| Ovarian Cancer | RMUGS | 0 | Atypical CNA profile |
| Ovarian Cancer | RMGI | 1 | Fairly quiet and atypical CNA profile; lacks TP 53 mutation and has several mutations and CNAs in cancer genes not typically altered in ovarian cancer |

## Discussion

While cell lines are valuable models of tumors, there are undeniable differences between the two ^2,,3,6-9,11-13,17,30-33^. Moreover, even tumors of a given cancer type/subtype may differ substantially from each other in terms of their genomic alterations ^34^. Thus, there is a need to refine comparisons of tumors samples, by focussing on the most important properties or alterations – at the same time, excluding all but the known important events means that we might miss potentially important shared alterations. Assigning weights to features of interest like specific genomic alterations, activation of a particular signalling pathway etc. allows us to incorporate the degree of importance of various features into our measure of similarity. Thus, including all or most available data along with judiciously chosen weights is an attractive option for comparing pairs of tumors and/or cell lines.

Here we introduced TumorComparer (TC), a weighted similarity based approach to assessing cell line – tumor similarity, and illustrated its use by comparing CCLE cell lines to TCGA tumors for six different cancer types. We used a set of weights – RA1 – that uses TCGA data to strongly emphasise cancer-type-specific recurrent genomic alterations, followed by pan-cancer recurrent alterations. We identified both good and poor genomic matches as well as outliers among the cell lines of all the cancer types. Several of the outliers and poor tumor matches were cell lines that lacked cancer-specific recurrent alterations reported by TCGA. We also flagged a few potentially mislabelled cell lines. Notably, while we found 13 outlier cell lines among ovarian cancer cell lines including five widely used ones, only 2-5 outliers were found in breast, colorectal, glioma, lung adenocarcinoma and lung squamous cell carcinoma cell lines (including no widely-used cell lines in lung adenocarcinoma, and only one each in the other cancer types). Thus the vast majority of cell lines, including most of the widely used ones, bear at least a moderate resemblance to tumors, in terms of sharing cancer-type-specific recurrent alterations, and not having an unusually high or low number of alterations. It is worth noting that the TCGA project has focussed on the genomics of primary, untreated tumors, and as such, is representative of that subset. Similarly, CCLE does not include all cell lines, and conclusions based on a comparison of TCGA and CCLE data may not apply to other collections of tumors and cell lines. However, the weighted similarity approach introduced here can be applied to genomic and molecular profiles beyond the TCGA and CCLE datasets analysed here.

Our method is widely applicable to comparisons of genomic profiles, including, but not limited to tumor-tumor, tumor–cell line and cell line–cell line comparisons. In particular, weighted similarity can be valuable for studies in precision medicine. For instance, given a set of patients with genomic profiles as well as treatment and outcome histories, we can compare a new patient’s genomic profile to the profiles of other patients using weighted similarity, and the history of the most similar patients can be used to gain insight into likely responses of the new patient to potential treatments. Similarly, if a set of patient data is not available, one can perhaps usefully exploit the availability of large drug sensitivity screens conducted in cell line panels ^16,35-38^ and make inferences based on cell lines most similar to a particular patient’s tumor regarding potential response to particular drug therapy.

This study has generalized our recent work on evaluating cell lines via comparison of genomic profiles in ovarian cancer ^17^, using weighted similarity with RA1, a set of weights chosen to emphasize important genomic alterations when computing pairwise similarity/distance. While the main conclusions of our previous study were reproduced by this more general approach, the assessment of individual cell lines (barring a few outliers) can vary, depending on the choice of similarity/distance measure, and the features we emphasize (and to what extent). In particular, the study on HGSOC (high-grade serous ovarian cancer) tumors and ovarian cell lines used TP53 mutation status, hypermutant status, and mutation status in seven “non-HGSOC” genes, along with correlation with the mean copy number alteration profile of tumors to score cell lines. The weighted similarity approach introduced here is more general and systematic, and all six cancer types/subtypes were studied using a consistent approach to deriving feature weights, and the same similarity measure. Our methodology can be applied to optimize comparison of cancer samples, be they *in-vivo* or *in-vitro,* in a flexible and data-driven manner. For instance, in cases where genomic similarity in terms of shared recurrent alterations might be deemed less important than other characteristics such as expression of specific biomarkers, or biological properties like growth characteristics or response to certain therapies^14^, our approach can be adapted to inform the comparison via the incorporation of features reflecting the characteristics of interest, and weights emphasizing said features. A particularly promising application is patient-patient similarity, which is going to be a critical component in personalized cancer therapy. As we acquire more molecular and clinical data along with treatment outcomes, meaningful measures of similarity to previously treated patients will be an invaluable guide for treatment strategies. By emphasising, via choice of weights, determinants of response and resistance to anti-cancer drugs, our approach can be adapted for use in prognosis, assignment to clinical trials and choice of therapy.

## Methods

### Data acquisition and preprocessing

TCGA data was obtained from the Broad Institute’s GDAC portal websites, and cell line data was obtained from the CCLE website ^16^. In order to focus on the mutations most likely to be functional, we excluded mutations in introns, 3’ and 5’ untranslated regions, flanking and intergenic regions, as well as silent and RNA mutations. Data was pre-processed using the Perl and R programming environments ^39^.

### GISTIC2 on CCLE CNA data

GISTIC2 ^18^ was run using the GenePattern ^40^ website, using the CCLE segmented data downloaded from the CCLE ^16^ website and all the default parameters, except the “confidence”, which was increased from 0.75 to 0.99. We used the discretized 5-valued (-2,-1,0,1,2) gene-wise data produced by the GISTIC algorithm for copy-number analysis.

### Assignment of weights to features

In general, feature weights are to be determined depending on the interest of the investigator and the question(s) asked. Here we chose a particular set of weights focussed on genomics alterations observed as recurrent across many cancer samples. We assigned each of the 3180 genomic features (mutations in 1651 genes, and copy number alterations (CNAs) in 1529 of these genes) a weight between 0 and 1 as follows (default weight = 0.01). As alterations in cancer genes that have no statistically significant recurrence in a particular cancer type may still be of biological interest, we gave all known cancer genes from the TCGA pan-cancer studies a weight of 0.1. Genes in the results from the recurrence analysis programs MUTSIG or GISTIC for specific tumor types have high weights as follows: (i) each gene which had a cancer-specific MUTSIG q-value ≤ 0.1 has a weight of 1; (ii) all genes in GISTIC peaks have a weight according to the number of genes in the peak - all genes in a peak that spans *n* genes have a weight of 1/*n*.

Thus, all genes with significant recurrence of mutation events according to the MUTSIG method and all genes in singleton peaks in the GISTIC method have the maximum possible weight of 1, other GISTIC peak genes have a weight inversely proportional to the size of their peak, all remaining known cancer genes have a weight of 0.1, and the remaining alterations, assumed to be passengers, have a weight of 0.01.

### Weighted asymmetric matching

The vast majority of features are zero in most samples, since only a small fraction of genes are mutated in a typical tumor – so 0-0 matches are the “default” or expected case, and not very informative. We computed the weighted similarity of two samples using weighted asymmetric matching, which measures the similarity between two samples after discarding the 0-0 matches (hence “asymmetric”, like the Jaccard Index for binary data).

Samples are represented by feature vectors

*X* = (*x*_1_, *x*_2_, …, *x_m_*) and *Y* = (*y*_1_, *y*_2_, …, *y_m_*) and a weight vector for feature weights *W* = (*w*_1_, *w*_2_, …, *w_m_*), their weighted similarity is calculated as

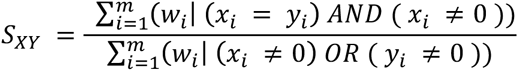

that is, 0-0 matches are discarded, and the similarity is calculated as the ratio of the sum of weights of features for which the two samples have the same value, to the sum of weights of all features for which at least one of the samples has a non-zero value. This is similar to the widely used Jaccard Index for binary data, in which zero-zero matches are discarded, and the similarity is calculated as the ratio of the intersection to the union of the subsets of features for which the two samples have non-zero values

### Evaluation of cell lines using MSK (mean similarity to k-nearest tumors) scores

Since tumor types and subtypes are often inherently heterogeneous, we evaluated cell lines using their similarity to a subset of *k* tumors, rather than all the tumors.

Once the weighted similarity scores have been computed, it is straightforward to determine the *k* most similar tumors for any given cell line or tumor (and *k*). Using the mean and standard deviation of the MSK scores of tumors, a cell line was deemed to be a poor match if its MSK score was more than two standard deviations below the mean MSK score for tumors, and an outlier if it was more than three standard deviations below the mean MSK score for tumors.

### Non-metric multidimensional scaling

We projected tumors and cell lines into two dimensions for approximate visual assessment of their proximity. Multidimensional scaling (MDS) is a method of dimension reduction which, given an input distance matrix, produces a mapping in a lower-dimensional space that preserves the distances in the original space as faithfully as possible. Classical MDS aims to directly compute the distances in the lower dimension so that they are as close to the original distances as possible (via a minimization of the sum of squares of error terms), and if used with Euclidean distances, is equivalent to PCA (principal component analysis) ^41^. Non-metric MDS, on the other hand, only aims to preserve the order between the distances, which allows it to potentially achieve a better low-dimensional mapping on datasets with a high variance in the distance matrix. Since only a low amount of the variance in our data was explained by the first two principal components, we used the *isoMDS* function from the R package *MASS* ^41^ to perform non-metric MDS. *isoMDS* uses the output of classical MDS (via the function *cmdscale*) as its initial configuration in the lower dimensional space, and then iteratively re-computes the distances in the lower dimensional space until convergence ^41^. Given weighted similarities between 0 and 1, distances were generated as

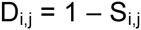

where D_i,j_ is the distance between samples i and j, and S_i,j_ is the similarity between them.

## Acknowledgements

We thank JianJiong Gao, Giovanni Ciriello, Yasin Senbabaoglou, Debora Marks and members of the Sander lab at MSKCC for discussions and Debra Bemis for discussions, manuscript edits and project management.

## Author Contributions

C.S. initiated and managed the project. R.S., C.S., and N.S. conceived the approach. R.S. developed and implemented the method, analysed the data and drafted the manuscript. R.S., C.S and N.S. interpreted the results and completed the manuscript.

## References

1 Masters, J. R. Human cancer cell lines: fact and fantasy. Nature reviews. Molecular cell biology 1, 233–236, doi:10.1038/35043102 (2000).

2 Wistuba, II et al. Comparison of features of human breast cancer cell lines and their corresponding tumors. Clinical cancer research: an official journal of the American Association for Cancer Research 4, 2931–2938 (1998).

3 Wistuba, II et al. Comparison of features of human lung cancer cell lines and their corresponding tumors. Clinical cancer research: an official journal of the American Association for Cancer Research 5, 991–1000 (1999).

4 Sandberg, R. & Ernberg, I. The molecular portrait of in vitro growth by meta-analysis of gene-expression profiles. Genome biology 6, R65, doi:10.1186/gb-2005-6-8-r65 (2005).

5 Ross, D. T. et al. Systematic variation in gene expression patterns in human cancer cell lines. Nature genetics 24, 227–235, doi:10.1038/73432 (2000).

6 Smiraglia, D. J. et al. Excessive CpG island hypermethylation in cancer cell lines versus primary human malignancies. Human molecular genetics 10, 1413–1419 (2001).

7 Houshdaran, S. et al. DNA methylation profiles of ovarian epithelial carcinoma tumors and cell lines. PloS one 5, e9359, doi:10.1371/journal.pone.0009359 (2010).

8 Hennessey, P. T. et al. Promoter methylation in head and neck squamous cell carcinoma cell lines is significantly different than methylation in primary tumors and xenografts. PloS one 6, e20584, doi:10.1371/journal.pone.0020584 (2011).

9 Tsuji, K. et al. Breast cancer cell lines carry cell line-specific genomic alterations that are distinct from aberrations in breast cancer tissues: comparison of the CGH profiles between cancer cell lines and primary cancer tissues. BMC cancer 10, 15, doi:10.1186/1471-2407-10-15 (2010).

10 Greshock, J. et al. Cancer cell lines as genetic models of their parent histology: analyses based on array comparative genomic hybridization. Cancer research 67, 3594–3600, doi:10.1158/0008-5472.CAN-06-3674 (2007).

11 Neve, R. M. et al. A collection of breast cancer cell lines for the study of functionally distinct cancer subtypes. Cancer cell 10, 515–527, doi:10.1016/j.ccr.2006.10.008 (2006).

12 van Staveren, W. C. et al. Human cancer cell lines: Experimental models for cancer cells in situ? For cancer stem cells? Biochimica et biophysica acta 1795, 92–103, doi:10.1016/j.bbcan.2008.12.004 (2009).

13 Kao, J. et al. Molecular profiling of breast cancer cell lines defines relevant tumor models and provides a resource for cancer gene discovery. PloS one 4, e6146, doi:10.1371/journal.pone.0006146 (2009).

14 Elias, K. M. et al. Beyond genomics: Critical evaluation of cell line utility for ovarian cancer research. Gynecologic oncology, doi:10.1016/j.ygyno.2015.08.017 (2015).

15 Collins, F. S. & Barker, A. D. Mapping the cancer genome. Pinpointing the genes involved in cancer will help chart a new course across the complex landscape of human malignancies. Scientific American 296, 50–57 (2007).

16 Barretina, J. et al. The Cancer Cell Line Encyclopedia enables predictive modelling of anticancer drug sensitivity. Nature 483, 603–607, doi:10.1038/nature11003 (2012).

17 Domcke, S., Sinha, R., Levine, D. A., Sander, C. & Schultz, N. Evaluating cell lines as tumour models by comparison of genomic profiles. Nature communications 4, 2126, doi:10.1038/ncomms3126 (2013).

18 Mermel, C. H. et al. GISTIC2.0 facilitates sensitive and confident localization of the targets of focal somatic copy-number alteration in human cancers. Genome biology 12, R41, doi:10.1186/gb-2011-12-4-r41 (2011).

19 Lawrence, M. S. et al. Mutational heterogeneity in cancer and the search for new cancer-associated genes. Nature 499, 214–218, doi:10.1038/nature12213 (2013).

20 Zack, T. I. et al. Pan-cancer patterns of somatic copy number alteration. Nature genetics 45, 1134–1140, doi:10.1038/ng.2760 (2013).

21 Lawrence, M. S. et al. Discovery and saturation analysis of cancer genes across 21 tumour types. Nature 505, 495–501, doi:10.1038/nature12912 (2014).

22 Kandoth, C. et al. Mutational landscape and significance across 12 major cancer types. Nature 502, 333–339, doi:10.1038/nature12634 (2013).

23 Cancer Genome Atlas Research Network. Integrated genomic analyses of ovarian carcinoma. Nature 474, 609–615, doi:10.1038/nature10166 (2011).

24 Mehta, J., O’Driscoll, L., Barron, N., Clynes, M. & Doolan, P. in Methods of Cancer Diagnosis, Therapy, and Prognosis Vol. 7 Methods of Cancer Diagnosis, Therapy and Prognosis (ed M. A. Hayat) Ch. 13, 183–191 (Springer Netherlands, 2010).

25 Chavez, K. J., Garimella, S. V. & Lipkowitz, S. Triple negative breast cancer cell lines: one tool in the search for better treatment of triple negative breast cancer. Breast disease 32, 35–48, doi:10.3233/BD-2010-0307 (2010).

26 Ciriello, G. et al. Emerging landscape of oncogenic signatures across human cancers. Nature genetics 45, 1127–1133, doi:10.1038/ng.2762 (2013).

27 Cancer Genome Atlas Research, N. Comprehensive genomic characterization defines human glioblastoma genes and core pathways. Nature 455, 1061–1068, doi:10.1038/nature07385 (2008).

28 Cancer Genome Atlas Research, N. Comprehensive molecular characterization of human colon and rectal cancer. Nature 487, 330–337, doi:10.1038/nature11252 (2012).

29 Mouradov, D. et al. Colorectal cancer cell lines are representative models of the main molecular subtypes of primary cancer. Cancer research 74, 3238–3247, doi:10.1158/0008-5472.CAN-14-0013 (2014).

30 Ertel, A., Verghese, A., Byers, S. W., Ochs, M. & Tozeren, A. Pathway-specific differences between tumor cell lines and normal and tumor tissue cells. Molecular cancer 5, 55, doi:10.1186/1476-4598-5-55 (2006).

31 Gillet, J. P. et al. Redefining the relevance of established cancer cell lines to the study of mechanisms of clinical anti-cancer drug resistance. Proceedings of the National Academy of Sciences of the United States of America 108, 18708–18713, doi:10.1073/pnas.1111840108 (2011).

32 Greshock, J. et al. Distinct patterns of DNA copy number alterations associate with BRAF mutations in melanomas and melanoma-derived cell lines. Genes, chromosomes & cancer 48, 419–428, doi:10.1002/gcc.20651 (2009).

33 Mehta, J. P., O’Driscoll, L., Barron, N., Clynes, M. & Doolan, P. A Microarray Approach to Translational Medicine in Breast Cancer: How Representative are Cell Line Models of Clinical Conditions? Anticancer Research 27, 1295–1300 (2007).

34 Wheeler, D. A. & Wang, L. From human genome to cancer genome: the first decade. Genome research 23, 1054–1062, doi:10.1101/gr.157602.113 (2013).

35 Garnett, M. J. et al. Systematic identification of genomic markers of drug sensitivity in cancer cells. Nature 483, 570–575, doi:10.1038/nature11005 (2012).

36 Shoemaker, R. H. The NCI60 human tumour cell line anticancer drug screen. Nature reviews. Cancer 6, 813–823, doi:10.1038/nrc1951 (2006).

37 Basu, A. et al. An interactive resource to identify cancer genetic and lineage dependencies targeted by small molecules. Cell 154, 1151–1161, doi:10.1016/j.cell.2013.08.003 (2013).

38 Klijn, C. et al. A comprehensive transcriptional portrait of human cancer cell lines. Nature biotechnology 33, 306–312, doi:10.1038/nbt.3080 (2015).

39 R: A Language and Environment for Statistical Computing (R Foundation for Statistical Computing, Vienna, Austria, 2015).

40 Reich, M. et al. GenePattern 2.0. Nature genetics 38, 500–501, doi:10.1038/ng0506-500 (2006).

41 Venables, W. N. & Ripley, B. D. Modern Applied Statistics with S. Fourth edn, (Springer, 2002).

